# Melanoma-specific mutation hotspots in distal, non-coding, promoter-interacting regions implicate novel candidate driver genes

**DOI:** 10.1101/2024.05.07.593069

**Authors:** Michael Pudjihartono, Nicholas Pudjihartono, Justin M. O’Sullivan, William Schierding

## Abstract

Investigations of the non-coding region are essential for uncovering the full spectrum of genetic drivers in melanoma. Here, we map distal, non-coding, promoter-interacting regulatory elements using Hi-C data from melanoma cells and integrating these with whole-genome sequence and gene expression data from the Pan Cancer Analysis of Whole Genomes. We identified eight recurrently mutated hotspots that are melanoma-specific, alter transcription factor binding motifs, and affect the expression of genes (e.g. *HSPB7*, *CLDN1, ADCY9* and *FDXR*) previously implicated as tumor suppressors/oncogenes. Our study integrates multiple levels of biological data to uncover novel melanoma-specific non-coding candidate drivers beyond the well-characterized *TERT* promoter.

## Introduction

Skin cancers rank as the most prevalent cancers worldwide^1^, with melanoma being the deadliest amongst them^2^. To develop targeted treatments, it is crucial to identify the entire spectrum of sites that, when mutated, are causative of melanoma development. However, a large fraction of mutations in cancer genomes are passengers, which do not confer a detectable selective advantage to cells and, therefore, do not undergo positive selection^3^. Conversely, regional recurrence of somatic mutations is indicative of the positive selection process that marks driver events, contributing to tumorigenesis^4^.

Large-scale cancer sequencing projects such as The Cancer Genome Atlas (TCGA)^5^ and the International Cancer Genome Consortium (ICGC)^6^ have initially focused on the coding regions, which led to the identification of hundreds of driver mutations in protein-coding genes^7^. In contrast, few non-coding drivers are currently known. Thus far, the most characterized non-coding driver is located in the telomerase reverse transcriptase gene (*TERT*) promoter, which upregulates *TERT* expression in melanoma and many other cancer types^8–10^. It is hypothesized that many other non-coding drivers exist outside the *TERT* promoter. And so, there has been a shift in interest to systematically analyze the non-coding regions—which represent the 98% of the human genome where the vast majority of somatic mutations reside^11^—for driver mutations^12^. However, most large-scale efforts have concentrated on gene-adjacent non-coding elements (e.g. promoters and non-coding RNAs^9,13–17^) leaving intergenic distal regulatory elements relatively unexplored.

Due to their role in regulating gene expression, distal regulatory elements represent a promising group of loci within the non-coding genome within which to investigate driver mutations. For example, recurrent mutations in distal regulatory elements of *PAX5*^18^ and *TAL1*^19^ have been identified in leukemia. However, these elements often regulate non-adjacent genes over large genomic distances^20–23^, making it challenging to systematically identify their true target genes based on linear genomic sequences. Hi-C technologies have emerged as valuable tools to identify long-range interactions within the 3D organization of the genome^24^. We have previously shown that such long-range connections can identify targets of germline gene regulation^25–27^. However, such genome organization varies across tissue types^28,29^, underscoring the need to analyze data from tissues relevant to the disease state.

This study leveraged melanoma-specific Hi-C data to map distal regulatory elements that physically interact with the promoter of 18,044 protein-coding genes. We integrate these with the TCGA/ICGC Pan-cancer Analysis of Whole Genomes (PCAWG) data on whole-genome sequence (WGS) and gene expression to identify eight candidate non-coding drivers. These drivers are novel, melanoma-specific, located in promoter-interacting distal regulatory elements, alter transcription factor binding motifs, and affect the expression of genes (e.g. *HSPB7*, *CLDN1, ADCY9* and *FDXR*) previously implicated as tumor suppressors/oncogenes in various cancers. In summary, our study integrates multiple levels of biological data to uncover melanoma-specific non-coding drivers, providing a solid foundation for targeted functional validations.

## Results

### Construction and characterization of a promoter-interaction network in melanoma

The hg38 human reference genome was *in-silico* digested at HindIII restriction enzyme cleavage sites (A/AGCTT), resulting in 851,637 non-overlapping genomic fragments [Figure 1a(i), Supplementary Table 1], with a median length 2219 bp [Supplementary Figure 1a]. A total of 41,733 unique genomic fragments overlapped a promoter region harboring active TSS^30^ [Figure 1a(ii), Supplementary Table 2], defined as “promoter fragments” [Supplementary Table 3].

**Figure 1.**
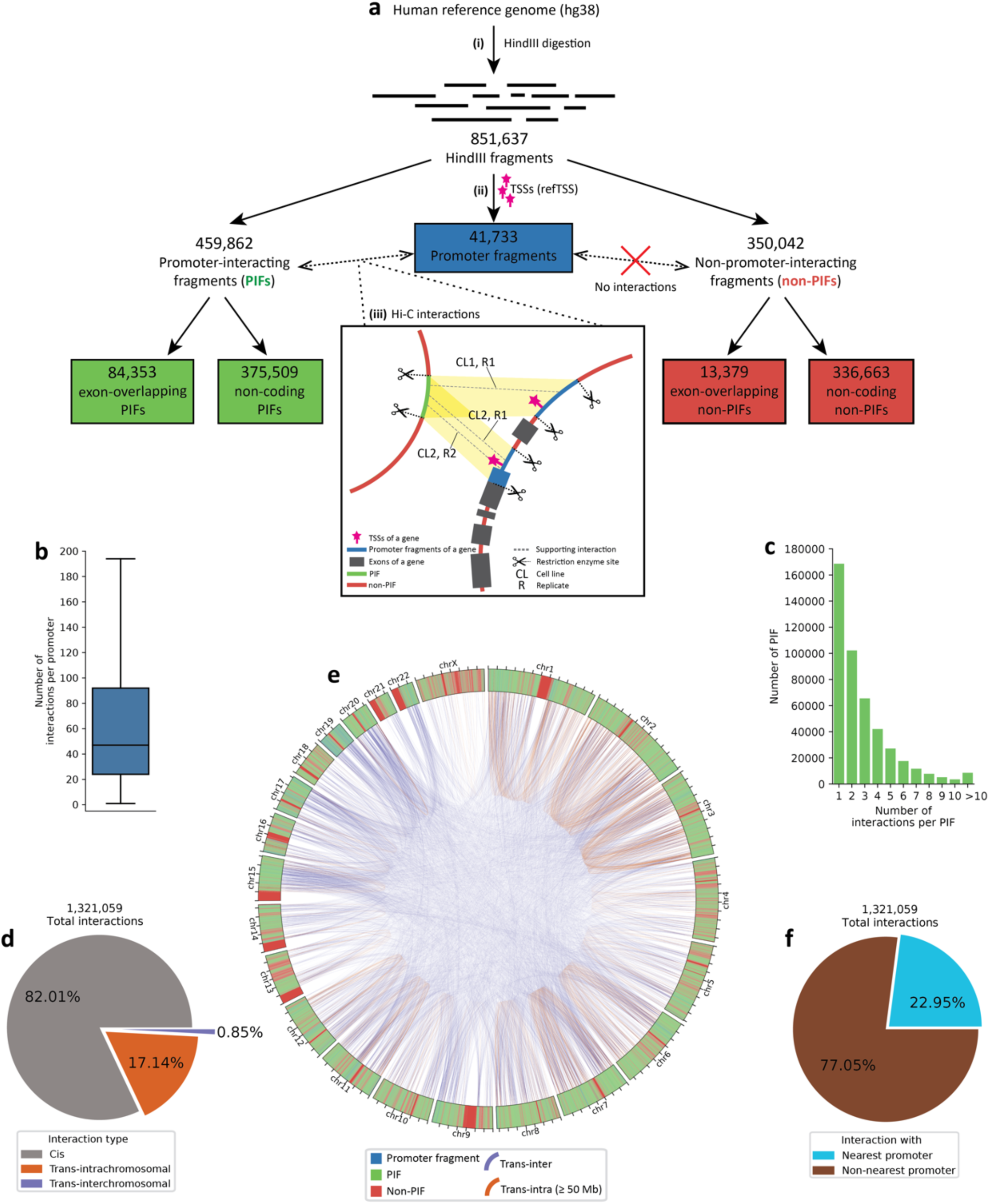
Workflow for identification of 375,509 non-coding, promoter-interacting fragments in melanoma-specific Hi-C cell-lines. (a) The workflow consisted of three parts: i) Digestion of the human genome into non-overlapping genomic fragments; ii) Identification of promoter fragments; iii) Identification of distal fragments interacting with the promoter fragment(s) of each protein-coding gene in the genome. In the cartoon example, a fragment-promoter interaction has a total of 3 supporting interactions from 3 different replicates of 2 different cell lines. The pipeline classified the total genomic fragments into distinct fragment classes (red, blue, and green boxes). (b) Number of interactions per promoter with distal promoter-interacting fragments (PIFs). (c) Number of interactions per PIF with promoters. (d) Proportion of different distance classes of interactions in relation to the total unique PIF interactions. (e) Circos plot illustrating the distribution of fragment classes across the genome and the interactions anchored at gene promoters. For clarity, only trans-inter and trans-intrachromosomal (≥ 50 Mb) interactions are shown (representing ∼1% of total interactions). (f) Proportions of interactions involving nearest promoters and those involving non-nearest promoters.

To catalog the putative regulatory elements specific to melanoma, we mapped all physical interactions between the promoter fragment(s) of each gene and the rest of the genome using Hi-C data from the two ENCODE melanoma cell lines (SK-MEL-5 and RPMI7951^31^) [Figure 1a(iii)]. This promoter-interaction network [Supplementary File 1] contained 459,862 unique non-promoter distal fragments that interacted with at least one gene promoter, referred to as “promoter-interacting fragments” (PIFs). The majority of these PIFs (n=375,509, 82%) were located within the non-coding regions of the genome (termed “non-coding PIFs”) [Figure 1a].

We observed that over 98% of gene promoters interacted with at least one PIF, with a median of 47 interactions per promoter [Figure 1b]. Conversely, most PIFs interacted with one or two gene promoters [Figure 1c]. These interactions primarily occurred within a distance of ≤ 1 Mb (*Cis*, 82% of the total interactions), while 17% occurred over distances > 1 Mb (*Trans-intrachromosomal*), with the remaining 1% involved interactions between chromosomes (*Trans-interchromosomal*) [Figure 1d]. The distribution of fragment classes across the genome and the interactions anchored at gene promoters is summarized in [Figure 1e].

We then investigated whether PIFs predominantly interacted with the nearest promoter. Our analysis revealed that 77% of PIF interactions occurred with a non-nearest promoter [Figure 1f]. This highlights the intricate and long-range spatial arrangements that potentially contribute to transcriptional regulation.

### Melanoma promoter-interacting fragments are under stronger purifying selection

To investigate the potential regulatory roles of the identified PIFs, we examined whether these PIFs showed evidence of purifying selection indicative of active regulatory sequences^32^. We employed the depletion rank (DR) score, a measure of genome-wide sequence conservation based on the UK biobank WGS data, where lower DR scores indicate regions under stronger sequence constraints^33^. Our analysis demonstrated that both exon-overlapping and non-coding PIFs exhibited significantly lower DR scores compared to their respective non-PIF counterparts [Figure 2a].

**Figure 2.**
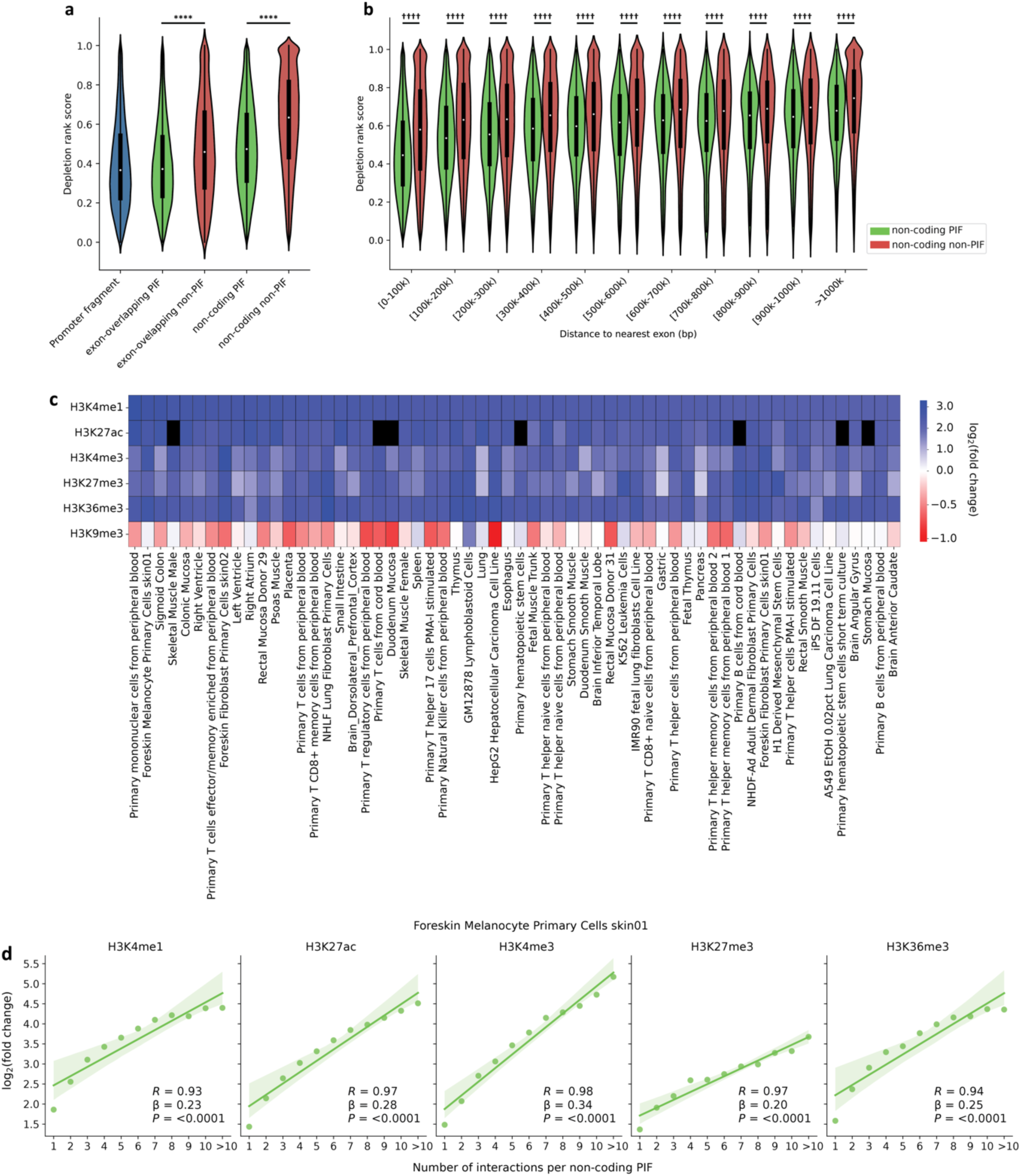
Promoter-interacting fragments (PIFs) are enriched for markers of active regulatory sequences. (a) Both exon-overlapping and non-coding PIFs are subject to increased purifying selection compared to their non-PIF counterparts, as shown by their significantly lower depletion rank (DR) score. Statistical significance was determined using the Wilcoxon rank-sum test. Significance levels: \**P* ≤ 0.05, \*\**P* ≤ 0.01, \*\*\**P* ≤ 0.001, \*\*\*\**P* ≤ 0.0001. (b) Non-coding PIFs consistently exhibited significantly lower DR scores than non-coding non-PIFs, regardless of their proximity to coding exons. Statistical significance was determined using the Wilcoxon rank-sum test with Bonferroni correction for multiple comparisons. Significance levels: †*adj. P* ≤ 0.05, ††*adj. P* ≤ 0.01, †††*adj. P* ≤ 0.001, ††††*adj. P* ≤ 0.0001. (c) Enrichment of non-coding PIFs for histone marks across tissues and cell lines. Cell lines are sorted based on the fold change of H3K4me1. The top 60 cell lines with the highest H3K4me1 fold change are displayed. Black cells represent cell lines with no available data in the Roadmap Epigenomics project. (d) For non-coding PIF, an increased number of promoter interactions correlates with higher levels of regulatory histone marks in melanocyte.

We observed that non-coding PIFs tended to be located in closer proximity to exons compared to non-coding non-PIFs [Supplementary Figure 1b]; at the same time, the DR score of fragments displayed an increasing trend with increasing distance from exons [Supplementary Figure 1c]. To ascertain whether the lower DR scores of non-coding PIFs relative to non-coding non-PIFs could be entirely attributed to the latter being situated further from exons, we evaluated the DR scores within distinct distance groups. Our findings indicated that distance from exon correlated with DR score for non-coding PIFs, but that distance was not the sole factor impacting the difference in DR score, as significant differences remained across every distance group [Figure 2b]. Therefore, the enrichment of lower DR scores in non-coding PIFs potentially reflects their regulatory characteristics, regardless of their proximity to coding exons.

### Melanoma promoter-interacting fragments are strongly enriched for skin-specific histone marks of gene regulation

Across 127 cell lines and tissues, non-coding PIFs were enriched in histone marks associated with enhancers and regulatory elements, with melanocyte and skin fibroblast tissues exhibiting strong enrichments [Figure 2c, Supplementary Table 4]. By contrast, H3K9me3—a histone mark associated with inactive heterochromatin^34^—was depleted across tissues. Additionally, a strong correlation was observed between the number of promoter interactions of non-coding PIFs and the enrichment of regulatory histone marks in melanocyte [Figure 2d]. Therefore, our finding is consistent with the potential transcriptional regulatory roles of the identified PIFs.

### Melanoma promoter-interacting fragments harbor melanoma-specific non-coding mutation hotspots

To investigate the impacts of somatic mutations within PIFs, we analyzed somatic single-nucleotide variations (SNVs) using WGS data from 107 melanoma patients in the PCAWG consortium. This cohort included 37 patients from the TCGA SKCM-US project and 70 from the ICGC MELA-AU project. Promoter fragments showed the lowest mutation rate among all fragment classes in this melanoma cohort [Figure 3a]. Importantly, exon-overlapping and non-coding PIFs exhibited significantly lower mutation rates compared to their respective non-PIF counterparts [Figure 3a]. This aligns with prior findings suggesting that functional regulatory regions tend to have reduced somatic mutation rates in cancers, possibly due to selective constraints or chromatin accessibility factors^17,35^.

**Figure 3.**
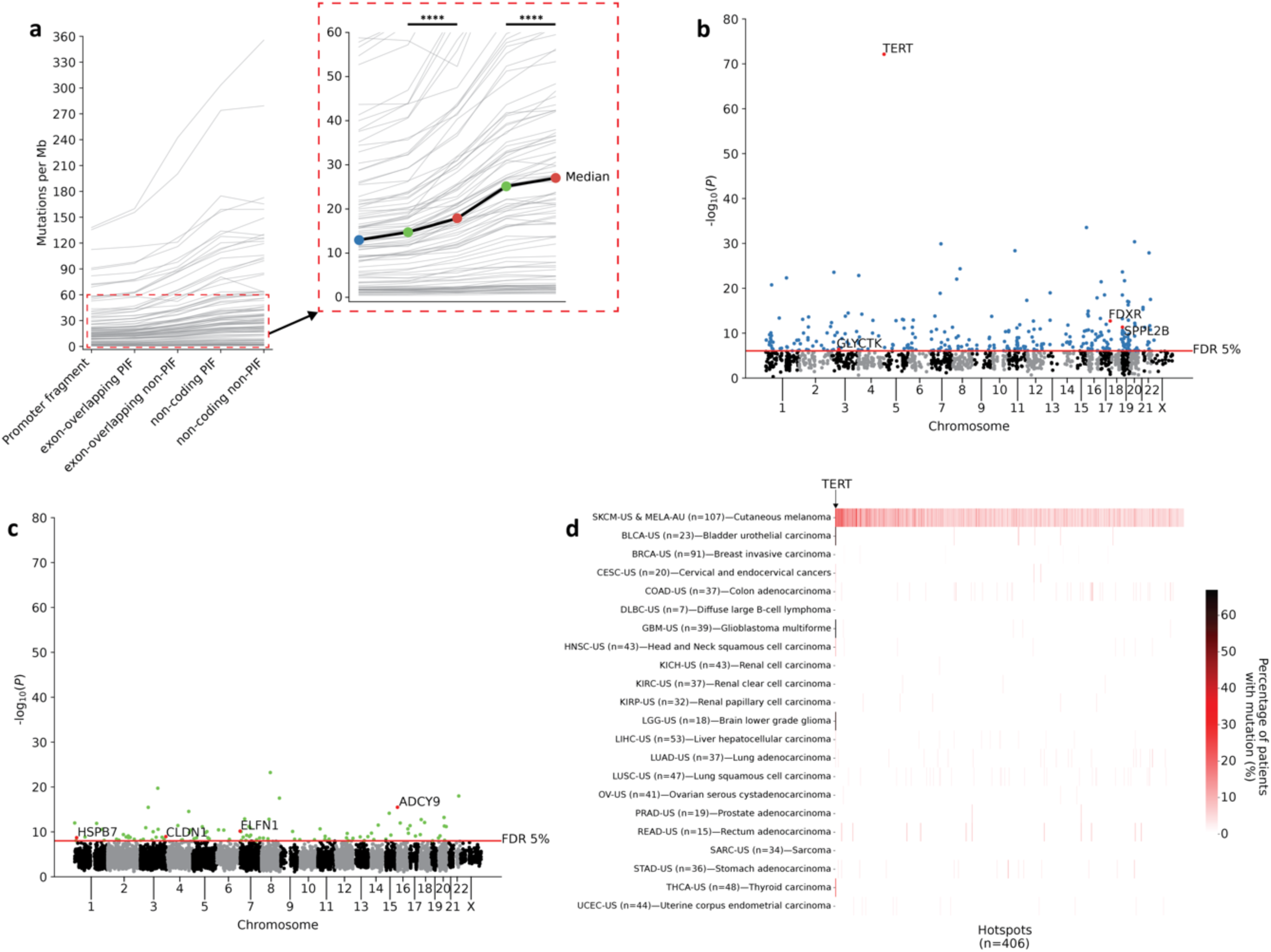
Non-coding somatic mutation hotspots in melanoma are melanoma-specific. (a) Promoter-interacting fragments (PIFs) have significantly depleted somatic mutation rates as compared to non-PIFs in the melanoma cohort of PCAWG. Grey lines represent data on individual patients (n = 107). The black line shows the median mutation rate across patients. Statistical significance was assessed using Wilcoxon signed-rank test. Significance levels: \**P* ≤ 0.05, \*\**P* ≤ 0.01, \*\*\**P* ≤ 0.001, \*\*\*\**P* ≤ 0.0001. (b and c) Manhattan plots of somatic mutation hotspots in (b) promoter fragments and (c) non-coding PIFs. Red dots indicate hotspots identified as significant (FDR ≤ 5%) somatic eQTLs in the melanoma-only and/or extended analyses [Figure 4a and b]. (d) Heatmap showing the percentage of patients (n=871) in each of the 22 cancer types with mutations in each melanoma hotspot (n = 406). Hotspots are sorted according to their *adj. P* values. Aside from *TERT*, most hotspots are melanoma-specific, leading to most areas of the plot having 0% non-melanoma patients with mutations (70% of the identified hotspots are exclusively mutated in melanoma alone).

The recurrence of somatic mutations above the expected background rate in specific genomic regions is suggestive of a positive selection process which may indicate driver events during tumor evolution^4^. To identify potentially functional non-coding somatic mutations, we utilized MutSpot^36^ to detect regions with such recurrence, termed “hotspots”, in promoter fragments and non-coding PIFs. To further account for melanoma-specific mutational biases^37,38^, we incorporated melanoma-specific epigenetic and sequence features alongside the standard MutSpot features into the selection of genomic features for the construction background mutation models (methods) [Supplementary Figure 2a and b].

Overall, the *TERT* promoter constituted the most significant hotspot in our analysis (*P* = 7.3 × 10^-73^) [Figure 3b]. However, beyond this well-documented hotspot^8–10^, we also identified 303 additional significant (FDR ≤ 0.05) hotspots in promoter fragments [Figure 3b, Supplementary Table 5a] and a further 102 in non-coding PIFs [Figure 3c, Supplementary Table 5b], totaling 406 significant hotspots. These hotspots are relatively short, with a median length of 41 bp and a maximum length of 114 bp. Of note, despite their lower mutation rate, promoter fragments harbored threefold more significant hotspots than non-coding PIFs and exhibit a lower *P* value overall [Figures 3b and c].

Cancer-associated germline single nucleotide polymorphisms (SNPs) identified by genome-wide association studies (GWAS) tend to reside in tissue or cancer type-specific regulatory elements^39–41^ and are implicated in modulating the activity of these elements^42^. To investigate whether the identified somatic mutation hotspots were pan-cancer or melanoma-specific, we analyzed the WGS data of an additional 764 patients across 21 cancer types from the TCGA portion of PCAWG (totaling 871 patients across 22 cancer types). Beyond a relatively small number of hotspots (*e.g.,* the *TERT* promoter, which is known to be highly mutated across cancer types^8–10^) the majority of the identified hotspots (n=281, 70%) are exclusively mutated in melanoma alone [Figure 3c]. Therefore, our approach captures signals of positive selection specific to melanoma.

### Melanoma-specific non-coding mutation hotspots alter the expression of eight candidate driver genes

The impact of a mutation on relevant biological functions can suggest its likelihood of being a driver^43^. Non-coding driver mutations should, therefore, act as distal regulators of gene expression. Changes at a locus that lead to changes in target gene expression are known as expression quantitative trait loci (eQTLs), which in germline contexts are often leveraged to infer the functional implications of non-coding GWAS risk SNPs^44^. We sought to determine whether the somatic mutations within melanoma-specific hotspots could influence tumor gene expression (termed “somatic eQTLs”).

Of the 107 patients in the melanoma PCAWG cohort with WGS data, 37 from the TCGA SKCM-US project had matching RNA-seq data available for this analysis. For each gene linked to at least one hotspot via our promoter-interaction network, we fit a multivariate linear regression model to evaluate its expression against the mutation status of its linked hotspot(s) and covariates such as copy number alteration, sex, and 7 hidden factors^45^ (methods) [Supplementary Table 6a]. This approach identified three hotspots as somatic eQTLs to three target genes at an FDR ≤ 0.05: *FDXR* and *GLYCTK* were significantly upregulated, while *ELFN1* was significantly downregulated [Figure 4a]. Relaxing the threshold to FDR ≤ 0.2 (as done in^46^) identified an additional suggestive association with the downregulation of *HSPB7* [Figure 4a].

**Figure 4.**
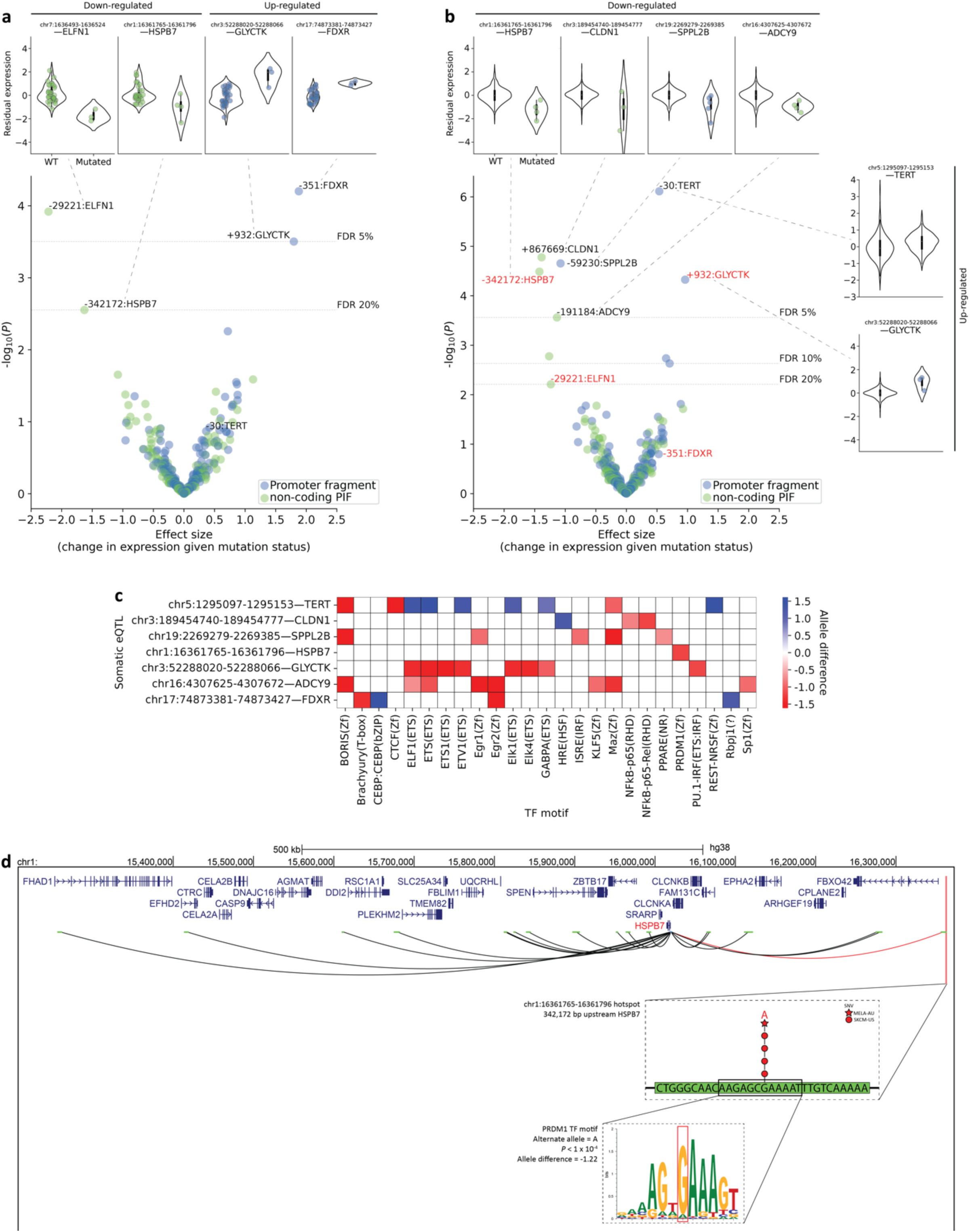
Eight melanoma somatic mutation hotspots are significantly associated with expression level changes of their Hi-C interacting target gene (FDR < 0.05, somatic eQTLs). (a) Melanoma-only analysis: three significant somatic eQTLs (FDR < 0.05), identified through multivariate linear regression, adjusting for copy number alteration, sex, and seven hidden factors. Violin plots detail the residual gene expression in wild-type (WT) vs. mutated patients for somatic eQTLs with FDR ≤ 20%. (b) Extended analysis (22 cancer types): six significant somatic eQTLs (FDR < 0.05), still targeting melanoma hotspots. This analysis incorporates additional covariates for cancer type, ancestry, and 16 hidden factors. Violin plots detail the residual gene expression in WT vs. mutated patients for somatic eQTLs with FDR ≤ 5%. (a and b) Blue and green colors in volcano plots indicate the fragment class encompassing each somatic eQTL. Somatic eQTLs in the volcano plots are labelled by the distance in base pairs relative to the start coordinate of the target gene in GENCODE v43 (adjusted for gene strand orientation). eQTLs with FDR ≤ 20% in (a) are highlighted red in (b) as comparison. (c) Prediction of transcription factor (TF) binding sites disrupted or created by SNVs in somatic eQTLs significant at FDR ≤ 5% in the melanoma-only and/or extended analyses. The DNA binding motif of each TF is listed in parenthesis on the x-axis. Allele difference is the difference between the TF binding score on the WT and mutated allele. Negative and positive values indicate the disruption and creation of TF motif, respectively. (d) Chromatin looping interactions between the *HSPB7* promoter and distal non-coding promoter-interacting fragments. A somatic eQTL is predicted to down-regulate *HSPB7* expression by disrupting a PRDM1 TF binding motif.

Given the constraints of our small cohort size and the potential for pan-cancer eQTL signals, a comparative re-analysis was conducted using data from all 22 TCGA cancer types within PCAWG (690 TCGA individuals with WGS and matching RNA-seq data; still targeting melanoma hotspots). Adjustments were made for additional covariates, including cancer type, ancestry, and 16 hidden factors (methods) [Supplementary Table 6b]. This extended analysis identified six somatic eQTL-target gene pairs at an FDR of ≤ 0.05, with the somatic eQTL for *TERT* showing the most significant association, consistent with the pan-cancer *TERT* signal found earlier^8–10^ [Figure 4b]. Apart from *TERT*, other somatic eQTL associations were melanoma-specific [Supplementary Table 6b]. The upregulated genes identified were *TERT* and *GLYCTK*, while the downregulated genes were *HSPB7*, *CLDN1*, *SPPL2B*, and *ADCY9*. Out of the four genes initially identified at an FDR ≤ 0.2 in the melanoma-only analysis, only *FDXR* falls below the FDR ≤ 0.2 cutoff in the extended analysis, and *HSPB7* reached FDR ≤ 0.05 [Figure 4b]. All four had the same direction of eQTL effect in both analyses.

One way that regulatory elements can modulate gene expression is by recruiting transcription factors (TFs) that bind to specific DNA sequence motifs^47^. Using motifbreakR^48^, we assessed the impact of SNVs within somatic eQTLs on TF binding motifs. Among the eight significant somatic eQTLs (FDR ≤ 0.05 in the melanoma-only and/or extended analyses), seven were found to harbor SNVs that disrupt or create at least one TF binding motif [Figure 4c, Supplementary Table 7]. Of the three genes with positive somatic eQTL effect sizes, two of them (*FDXR, TERT*) also show positive motif binding (motif creation), while only one of the four down-regulated genes had a positive motif binding score. For example, the G>A mutation within the eQTL at chr1:16361765-16361796 is predicted to strongly disrupt the binding affinity of the PRDM1 TF, thus potentially explaining its association with the downregulation of *HSPB7*, located 342,173 bp away [Figure 4d]. Interestingly, this somatic eQTL showed no association with its nearest expressed promoter, *FBXO42*, just 9,286 bp away [Supplementary Figure 3a]. *ELFN1* was the only significant somatic eQTL at a non-coding PIF to associate with its nearest promoter [Supplementary Figure 3a-c].

## Discussion

Identifying the full spectrum of genetic alterations driving melanoma is fundamental to understanding the molecular etiology of melanoma and progressing towards more advanced, targeted therapies. Given the potential of non-coding mutations in altering gene regulatory mechanisms and melanoma development, our study aims to uncover hidden drivers in these regions. By leveraging melanoma-specific Hi-C data, we identified distal non-coding genomic fragments that interact physically with well-annotated promoters of 18,044 protein-coding genes. These fragments, termed “non-coding promoter-interacting fragments” (non-coding PIFs), form the basis of our approach, which, while conceptually similar to promoter-capture Hi-C (PCHi-C), leverages available Hi-C data to provide a viable framework for promoter-interaction analysis in the absence of melanoma-specific PCHi-C datasets. The non-coding PIFs are under stronger sequence constraint, enriched for histone marks of gene regulation, and harbor lower somatic mutation rate compared to their non-promoter-interacting counterparts (non-coding non-PIFs), making them suitable candidate regions for investigating driver mutations.

Using WGS and RNA-seq data from the PCAWG project, we discovered 102 and 304 hotspots harboring recurrent SNVs in non-coding PIFs and promoters, respectively, within melanoma samples. Among these, 3 hotspots—1 in non-coding PIF and 2 in promoters—were identified as somatic eQTLs affecting target gene expressions in melanoma [Figure 4a]. Extending the analysis of the same hotspots across all 690 TCGA cancer patients within PCAWG identified 5 additional somatic eQTL associations. This suggests that the influence of many other hotspots may extend beyond transcriptional regulation, or that their effects on gene expression are subtler than detectable with our current statistical power. Additionally, despite our thorough adjustments for established general and melanoma-specific mutation rate covariates, it remains possible that some of the non-eQTL hotspots may be passenger or false positives.

Our study identified a total of eight target genes whose expression levels are altered by eight somatic eQTLs, seven of which contain SNVs that interfere with TF binding motifs. Aside from *TERT*, the other seven eQTLs are melanoma-specific, and their target genes have not been appreciably recognized as melanoma drivers. Nonetheless, most have been implicated as tumor suppressors or oncogenes in various cancer types. For example, *HSPB7*, a member of the small heat shock protein family, has been implicated as a tumor suppressor in renal cell carcinoma (RCC), where it is commonly downregulated^49^. Consequently, ectopic introduction of *HSPB7* suppressed RCC cancer cell lines growth^49^. Here, we found *HSPB7* as being downregulated by a somatic eQTL at non-coding PIF positioned 342,173 bp upstream. Furthermore, this somatic eQTL contain SNVs that strongly disrupt the binding site of PRDM1 TF. *PRDM1* is a tumor suppressor in melanoma^50^ and other cancers^51–54^, and loss of *PRDM1* has been shown to accelerate the onset and progression of melanoma^50^.

Additionally, we identified *CLDN1* as being downregulated by a somatic eQTL in non-coding PIF positioned 867,699 bp downstream. Known for its role in cell adhesion, *CLDN1* is downregulated in metastatic melanomas compared to benign nevi^55^. The introduction of *CLDN1* into melanoma brain metastatic cells expressing low levels of *CLDN1* has been shown to suppress their metastatic phenotype^55^. *CLDN1* is also implicated as a tumor suppressor in lung adenocarcinoma^56^, prostate^57^ and ER+ breast cancer^58^. Similarly, *ADCY9*, downregulated by a somatic eQTL in non-coding PIF positioned 191,184 bp upstream, has been recently identified as a novel tumor suppressor in lung adenocarcinoma^59^. Furthermore, we discovered a somatic eQTL in the *FDXR* promoter, associated with *FDXR* upregulation. Supporting this, a recent study has shown that *FDXR* drives the proliferation of primary and endocrine-resistant breast cancer cells by promoting mitochondrial fatty acid oxidation through *CPT1A* upregulation^60^. Importantly, *CPT1A* is a potential therapeutic target in melanoma, with its knockdown shown to inhibit the proliferation of V600E melanoma cells^61^. Thus, our study identified seven novel genes potentially driving melanoma, inviting deeper investigations to clarify their roles in melanoma.

Long-range *trans* associations are pivotal in elucidating the role of germline GWAS SNPs in complex diseases^26,62–64^, suggesting their potential significance in elucidating the impacts of somatic mutations in cancer. Challenges such as limited sample sizes have constrained the study of germline *trans*-eQTLs^65^. Given the power of detection and low sample sizes, our somatic eQTL analyses were only powered to detect *cis*-eQTLs. Therefore, all significant eQTL associations corresponded to *cis*-regulated genes. However, most significant somatic eQTLs at non-coding PIFs (3/4) targeted genes located beyond 100,000 bp away and lacked associations with their nearest promoters. The most distant association was observed for *CLDN1*, with its regulating somatic eQTL positioned 867,669 bp downstream. Additionally, *MSR1* (a tumor suppressor in prostate cancer^66^ and chronic myeloid leukemia^67^) presented the most suggestive *trans* association, downregulated by a somatic eQTL positioned 1.5 Mb downstream (*P*=0.017, FDR≤0.4). These observations highlight the limitation of the nearest-gene approach in identifying true target genes, which overlooks the complexity of long-range gene regulatory interactions. Our approach, which identifies somatic eQTL associations within hotspots with confirmed physical interactions in melanoma cell lines, not only improves the accuracy in pinpointing their true tissue-specific target genes but also narrows down the search space, thus enhancing the power to detect somatic eQTLs.

The datasets used in this study present both strengths and limitations. Our use of melanoma-specific Hi-C datasets to construct a promoter-interaction network captures the tissue-specificity of 3D genome organization^28^. Nonetheless, our current datasets do not address the dynamic^68^ and potentially individual-specific nature of the 3D genome organization, which extends beyond tissue-specificity. Furthermore, Hi-C, unlike PCHi-C, isn’t optimized for detecting promoter interactions and thus may not capture all interactions due to its lower effective sequence depth for these regions^69,70^. Therefore, our analysis may overlook additional long-range interactions that a melanoma-specific PCHi-C could potentially reveal. Finally, our analysis is constrained by the limited expression data for melanoma, with only 37 matched WGS and RNA-seq datasets available. Despite these limitations, we have identified several melanoma-specific hotspots of somatic mutations affecting target gene expressions and anticipate discovering more as additional matched datasets become available.

## Conclusion

In conclusion, our study suggests additional non-coding drivers beyond the well-characterized *TERT* promoter in melanoma. These drivers are melanoma-specific, located in promoter-interacting DNA regions, alter transcription factor binding motifs, and affect the expression of genes previously implicated as tumor suppressors/oncogenes in various cancer types. More broadly, our study provides a framework for integrating multiple levels of biological data to uncover cancer-specific non-coding drivers, providing a solid foundation for targeted functional validations.

## Materials and Methods

### In-silico digestion of the human genome

The hg38 human reference genome sequence was downloaded from NCBI (https://www.ncbi.nlm.nih.gov/datasets/genome/GCF_000001405.26/) and digested *in-silico* at HindIII restriction enzyme cleavage sites (A/AGCTT) into 851,637 non-overlapping genomic fragments using the Restriction package of Biopython^71^.

### Defining gene promoters

Data on active transcription start sites (TSSs) (i.e. refTSS_v3.3_human_coordinate.hg38.bed and refTSS_v3.3_human_annotation.txt) was obtained from refTSS v3.3^30^ (https://reftss.riken.jp/datafiles/ <u>3.3/human/</u>). Each TSS was mapped to one protein-coding gene annotated by GENCODE v43^72^, and TSSs located on chrM and chrY were excluded.

For each TSS mapped to the same gene, a 2,000 bp upstream and 200 bp downstream extension (relative to the DNA strand of the mapped gene) was performed, followed by merging of any overlapping intervals. The resulting interval(s) define the gene’s promoter region(s). The pybedtools^73^ python library was used to identify the genomic fragment(s) encompassing the gene’s promoter region(s), thus defining the “promoter fragment(s)” of that gene. In total, we identified the promoter fragments of 18,044 protein-coding genes located within 41,733 unique genomic fragments.

### Construction of a melanoma promoter-interaction network

As the spatial interactions of genomes are tissue-specific^28,29^, we focused on analyzing Hi-C chromatin interaction libraries from two melanoma cell lines: SK-MEL-5 and RPMI-7951 (two replicates each). Raw Hi-C data were downloaded from GEO (https://www.ncbi.nlm.nih.gov/geo/, accession: GSE105491 and GSE106022 for SK-MEL-5 and RPMI-7951, respectively) and processed (as previously described^74^) to obtain Hi-C chromatin interaction library files with the following format: read name, strand1, chr1, position1, fragment1, strand2, chr2, position2, fragment2. Each row in the processed interaction files describes the alignment of the two interacting read pairs (i.e. 1 and 2). Our pipeline works with any Hi-C pipeline that can generate interaction files in the required data format (e.g. HOMER^75^, Juicer^76^).

To characterize the regulatory landscape of melanoma, we mapped the genome-wide physical interactions anchored to the promoter fragment(s) identified for each gene. These promoter interactions were identified on a presence/absence basis. A fragment-promoter interaction is valid whenever it has ≥2 supporting interactions from ≥2 different replicates of ≥1 cell lines. The resulting promoter-interaction network [Supplementary File 1] identified 459,862 non-promoter distal fragments (promoter-interacting fragments; PIFs) that interacted with at least one gene promoter, making 1,321,059 unique interactions.

Overall, this approach allowed for the classification of the total genomic fragments (n=851,637) into distinct fragment classes:

1. Promoter fragment (n=41,733)
2. PIF: exon-overlapping (n=84,353) or non-coding (n=375,509)
3. Non-PIF: exon-overlapping (n=13,379) or non-coding (n=336,663)

### Sequence constraint analysis

We downloaded the raw depletion rank (DR) score data from the supplementary data of Halldorsson et al^33^. This dataset (i.e. DR.gor) contains the associated DR score for each overlapping 500-bp windows in the genome with a 50-bp step size. The pybedtools Python library was used to identify every 500-bp window that had an overlap of more than 50% with each of the 851,637 genomic fragment. The median DR score for each fragment was then calculated and assigned as the corresponding DR score for that fragment. Statistical significance was determined using Wilcoxon rank-sum test with Bonferroni correction for multiple comparisons.

### Histone mark enrichment analysis

We downloaded narrowpeak ChIP-seq data on 6 histone marks (H3K4me1, H3K27ac, H3K4me3, H3K27me3, H3K36me3 and H3K9me3) from 127 tissues and cell lines from Roadmap Epigenomics^77^ (https://egg2.wustl.edu/roadmap/data/byFileType/peaks/consolidated/narrowPeak/). All genomic coordinates were converted to hg38 using LiftOver^78^ tool. These data were then overlapped with the non-coding PIFs that we identified. We selected random non-coding genomic fragments that matched the total length of the identified non-coding PIFs. We performed resampling of the random non-coding genomic fragments 1,000 times and assessed the statistical significance of the enrichment using permutation test. Fold changes were calculated by dividing the rate of overlap of ChIP-seq peaks in non-coding PIFs and in non-coding non-PIFs, and subsequently, log_2_ transformed.

### Identification of somatic mutation hotspots

The PCAWG SNV/INDEL concensus callsets of ICGC (*i.e.*, final_consensus_snv_indel_passonly_ icgc.open.tgz) and TCGA (*i.e.*, final_consensus_snv_indel_tcga.controlled.tgz) were downloaded from the ICGC portal (https://dcc.icgc.org/releases/PCAWG/consensus_snv_indel/). All genomic coordinates were converted to hg38 using LiftOver tool. MutSpot^36^ was then used to identify focal genomic regions (hotspots) harboring recurrent SNVs identified from 70 ICGC and 37 TCGA melanoma patients. MutSpot was run separately on promoter fragments and non-coding PIFs by specifying the *region.of.interest* option. Additionally, the following options were specified:

1. The LASSO stability threshold, *cutoff.nucleotide* and *cutoff.features*, was set at 1 and 0.98, respectively.
2. The window size for hotspot discovery, *hotspot.size*, was set to 31bp.
3. Minimum number of mutated samples in each hotspot, *min.count*, was set to 4.

MutSpot provides a default pool of 135 features as potential covariates in the background mutation model calculation. As suggested^36^, we added the following additional features to correct for known melanoma-specific mutational biases:

1. Narrowpeak ChIP-seq on 6 histone marks (H3K4me1, H3K27ac, H3K4me3, H3K27me3, H3K36me3 and H3K9me3) and DNase-seq data from melanocytes to correct for melanocyte-specific local variations in mutation rate (downloaded from Roadmap Epigenomics as described earlier).
2. Intervals of CTTCCG motif to correct for known vulnerability to UV-induced mutagenesis at these sites^37^.
3. Intervals of transcription factor binding sites in melanoma cell line to correct for known hypermutations due to impaired nucleotide excision repair at these sites^38^ (downloaded from http://bg.upf.edu/group/projects/tfbs/).

Significant hotspots were determined using an FDR cutoff of ≤ 0.05. Hotspots at fragment boundaries were excluded from further analyses.

### Somatic expression quantitative trait loci (eQTL) analysis

For the somatic eQTL analyses, only hotspots that are mutated in ≥ 3 SKCM-US patients were considered. For each gene linked to at least one of these hotspots via our promoter-interaction network, we fit a multivariate linear regression model to evaluate its expression against the mutation status of its linked hotspot(s) and other covariates as follows:

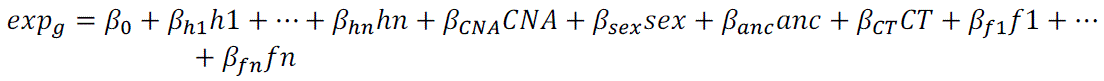

Where *exp*_g_ represent the normalized gene expression level *exp* of gene *g* and β values represent the regression coefficients for:

- The mutation status of its linked hotspot(s), ℎ (0=wild-type, 1=mutated)
- Copy number alteration levels, *CNA* (0=neutral, 1=amplified, 2=highly amplified, –1=deleted, –2=deeply deleted)
- Sex, *sex* (0=female, 1=male)
- Ancestry, *anc* (5 ancestries: European/African/East Asian/Admixed American/South Asian, one-hot encoded)
- Cancer type, *CT* (22 cancer types, one-hot encoded)
- Hidden factors, *f* (real values)

For the melanoma-only analysis, the covariates considered were ℎ, *sex*, *CNA*, and *f* because this cohort is homogeneous with respect to *CT* and consisted exclusively of individuals of European *anc*. The extended analysis, however, incorporated all covariates. Information on *sex*, *anc*, and *CT* for each patient was obtained from Supplementary Table 1 of the main PCAWG publication^79^.

Normalized RSEM RNA-seq data (i.e. *.rnaseqv2 illuminahiseq_rnaseqv2 unc_edu Level_3 RSEM_genes_normalized data.data.txt) were downloaded from GDAC firehose (https://gdac.broadinstitute.org). This normalization involved dividing the raw RSEM counts of each gene by the 75^th^ percentile value within the corresponding patient and multiplying by a factor of 1000. Genes were selected for further analysis if their normalized RSEM values exhibited a median of > 1 across all patients, resulting in 16,109 and 16,604 expressed genes in the melanoma-only and extended analysis, respectively. Normalized RSEM values were further log_2_ transformed and z-score standardized.

Hidden factors *f*s influencing gene expression were identified using probabilistic estimation of expression residuals (PEER)^45^. For the melanoma-specific and extended analyses, we estimated 20 and 50 potential *f*s, respectively, incorporating a mean term and covariates *sex*, *anc*, and *CT* during estimation. Analyses of the posterior variance of factor weights suggested 7 and 16 optimal *f*s for the melanoma-only and extended analysis, respectively.

PCAWG consensus callset for *CNA* (i.e. all_samples.consensus_CN.by_gene.170214.txt) was downloaded from the ICGC portal (https://dcc.icgc.org/releases/PCAWG/consensus_cnv/gene_level_ <u>calls</u>). Genes with missing *CNA* data in at least one patient were removed from further analysis, resulting in 23,171 genes with *CNA* data. Ultimately, 291 and 298 genes linked to at least one ℎ, expressed, and with *CNA* data were analyzed in the melanoma-only and extended analysis, respectively.

For each gene, feature selection was achieved by adding an L1-norm to the objective function, which minimizes the squared error between actual and predicted gene expression levels:

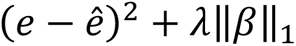

The regularization parameter λ was optimized through 10-fold cross-validation. For instances where the L1 penalty led to all β_ℎ_ coefficients being zero, λ was decreased to ensure inclusion of at least one hotspot. After fitting the multivariate linear regression model with the optimal parameter sets, we derived a gene-level and individual hotspot-gene *P* values (some genes are linked to >1 hotspots). For the gene-level *P* value, we used F-statistics by comparing the full model accuracy against that of a simpler model without any hotspots. Individual hotspot-gene *P* value and effect size was obtained directly from the regression model. Multiple hypothesis was corrected using the Benjamini-Hochberg method independently for gene-level and individual hotspot-gene *P* values. This overall approach for identifying somatic eQTLs is similar to what have been previously implemented by Zhang et al.^46^ and Soltis et al.^80^.

### Transcription factor (TF) binding analysis

We used motifbreakR (v2.16.0)^48^ to predict the impact of SNVs on TF bindings, utilizing position weight matrices of 247 human TFs from the HOMER^75^ motif data source provided by MotifDb (v1.42.0)^81^. This analysis focused on SNVs within the eight significant (FDR ≤ 0.05) somatic eQTLs in the melanoma-only and/or extended analyses. We used the “information content” scoring algorithm and applied a threshold of *P* < 1 x 10^-4^ to determine a motif match. motifbreakR classifies the normalized binding score difference between the reference and the alternate allele of each SNV as “neutral” (allele difference < 0.4), “weak” (allele difference < 0.7), or “strong” (allele difference ≥ 0.7). We only consider strongly affected motifs.

## Supplementary Figures

**Supplementary Figure 1.**
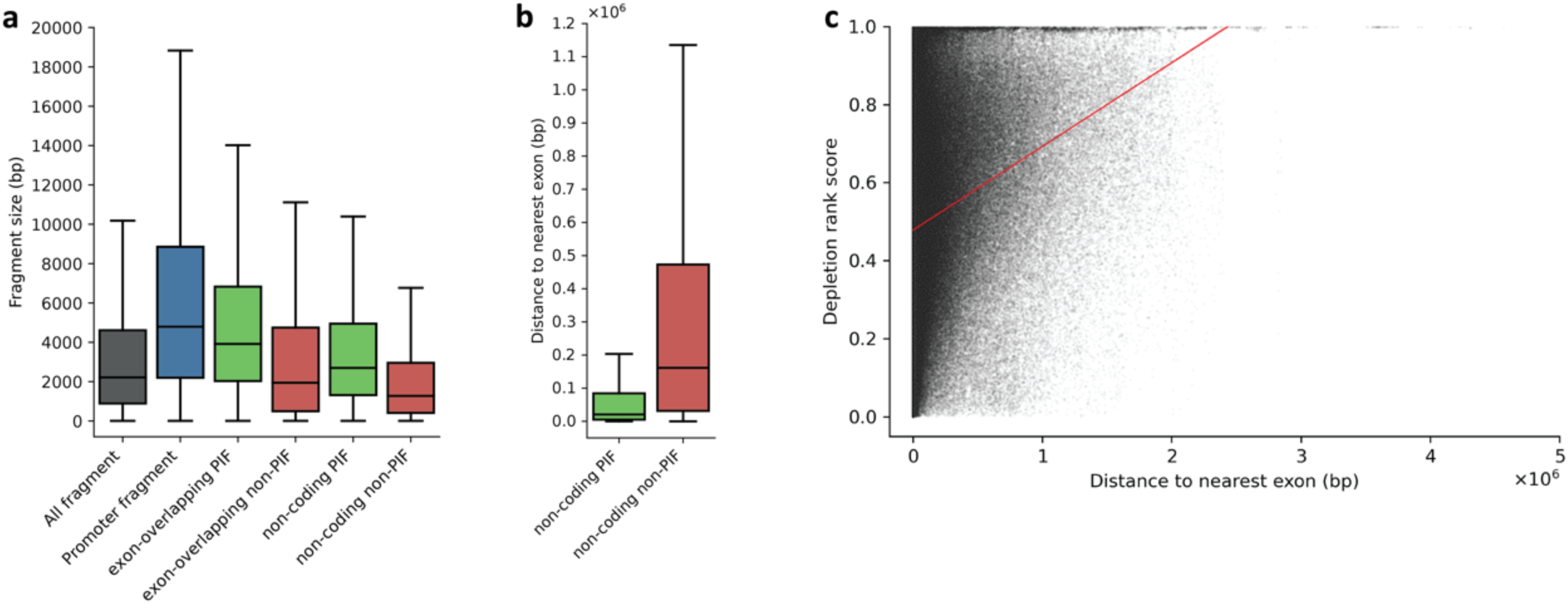
Characteristics of genomic fragment classes. (a) Distribution of fragment sizes across different fragment classes. (b) non-coding promoter-interacting fragments (PIFs) are located in closer proximity to coding exons compared to non-coding non-PIFs. (c) Depletion rank score of each individual fragment plotted against the distance to the nearest coding exon. Red line indicates the linear regression line of best fit.

**Supplementary Figure 2.**
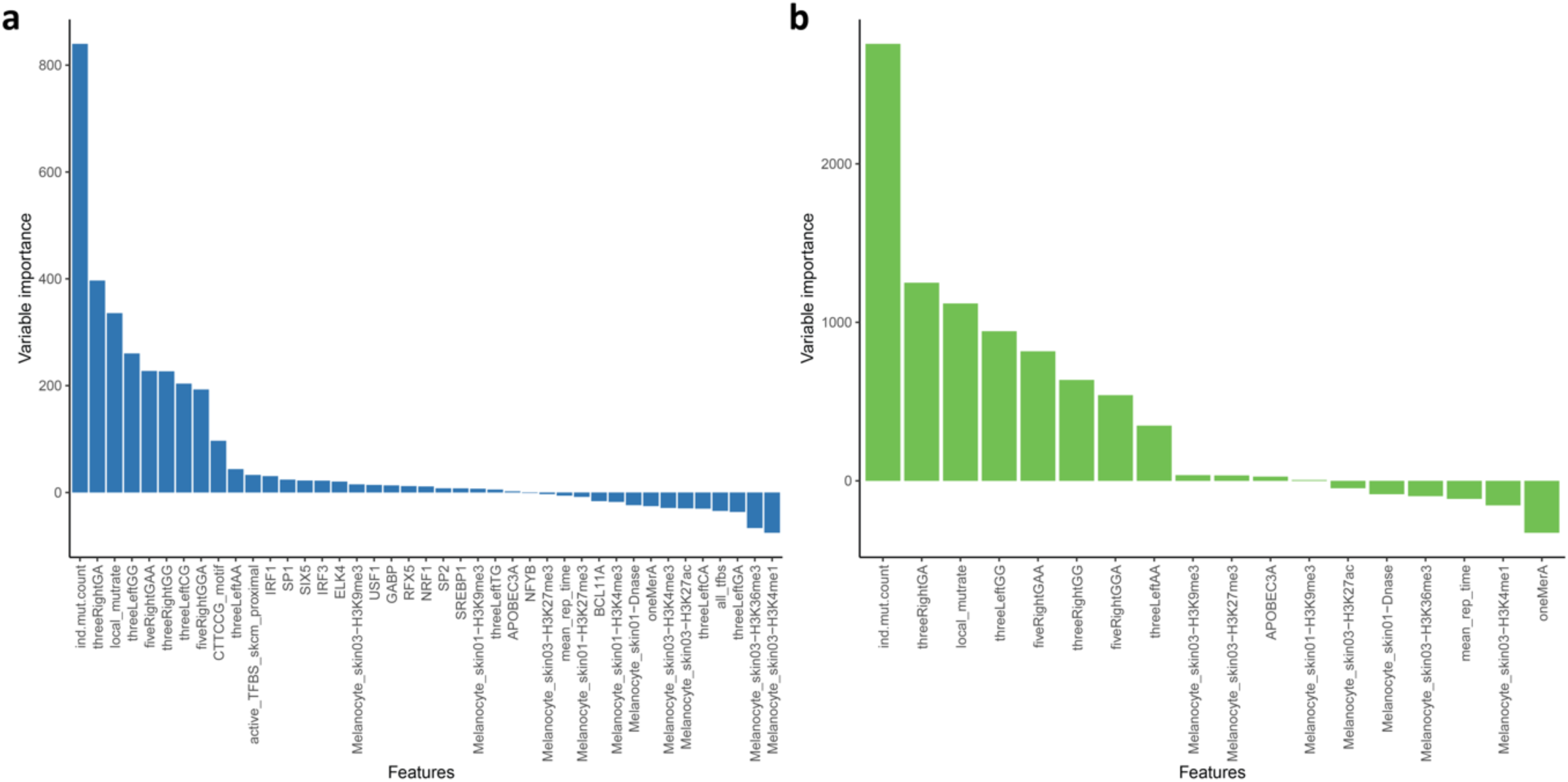
Bar plot of feature importance of the background mutation model outputted by MutSpot for (a) promoter fragments and (b) non-coding promoter-interacting fragments.

**Supplementary Figure 3.**
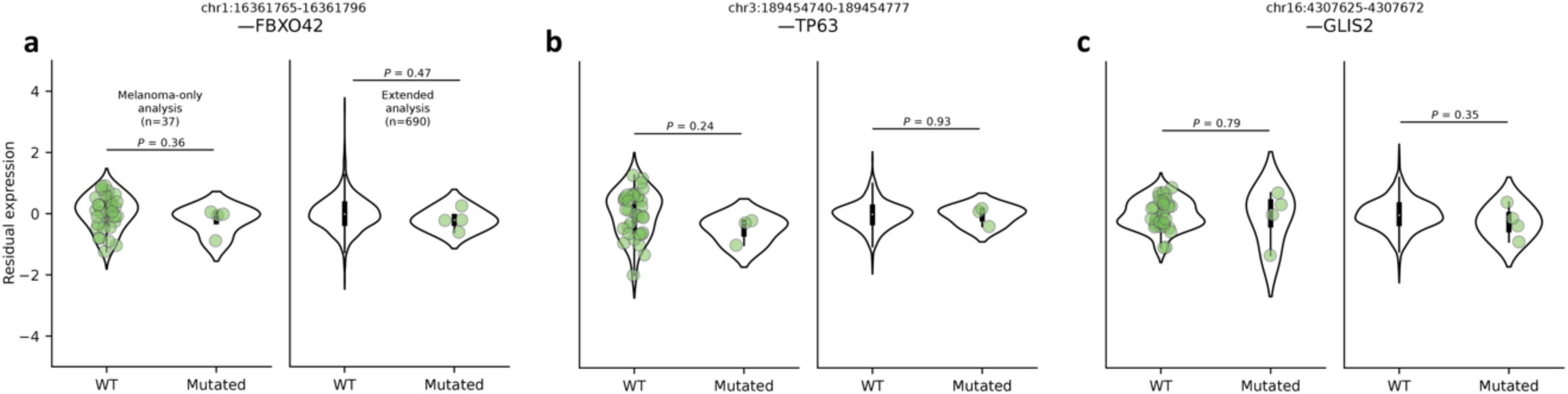
Significant somatic eQTLs at non-coding promoter-interacting fragments lacked associations with their nearest gene promoters. The violin plots compare residual gene expression levels between wild-type (WT) and mutated patients, specifically highlighting the hotspots (a) chr1:16361765-16361796 (somatic eQTL to *HSPB7*) and its nearest promoter, (b) chr3:189454740-189454777 (somatic eQTL to *CLDN1*) and its nearest promoter, and chr16:4307625-4307672 (somatic eQTL to *ADCY9*) and its nearest promoter.

## Acknowledgement

JOS was funded by donations from the Dines Family trust. WS was supported by a postdoctoral fellowship from the Vision Research Foundation and a Royal Society of New Zealand Marsden Grant (20-UOA-002). MP and NP were funded by a University of Auckland doctoral scholarship.

## Author contributions

MP contributed to conceptualization, performed analyses, data interpretation, and wrote the manuscript. WS and JOS supervised MP, conceptualized, and co-wrote the manuscript. NP contributed to data processing for somatic eQTL analyses and revisions of the manuscript. All authors contributed to the article and approved the submitted version.

## Data and code availability

Access to the controlled SNV/INDEL consensus callsets of TCGA was approved for General Reseach Use by the dbGaP (https://www.ncbi.nlm.nih.gov/gap/) Data Access Committee (Project ID: 34995, accession: phs000178). The “Promoter Interaction Network” software used to construct and characterize melanoma promoter-interaction network is available on GitHub (https://github.com/ <u>MichaelPudjihartono/Promoter-Interaction-Network</u>). The “regionperm” software used to assess the significance of histone mark enrichments through permutation tests is available on GitHub (https://github.com/MichaelPudjihartono/regionperm). Data analyses and visualizations were performed using Python (version 3.8.12) through Jupyter notebook (version 6.4.6) or using R (version 4.0.4) through RStudio (version 1.4.1106). Additional in-house scripts used for data wrangling are available upon request.

